# Automatic annotation of Cryo-EM maps with the convolutional neural network Haruspex

**DOI:** 10.1101/644476

**Authors:** Philipp Mostosi, Hermann Schindelin, Philip Kollmannsberger, Andrea Thorn

## Abstract

In recent years, three-dimensional density maps reconstructed from single particle images obtained by electron cryo-microscopy (Cryo-EM) have reached unprecedented resolution. However, map interpretation can be challenging, in particular if the constituting structures require de-novo model building or are very mobile. Here, we demonstrate the potential of convolutional neural networks for the annotation of Cryo-EM maps: our network *Haruspex* has been trained on a carefully curated set of 293 experimentally derived reconstruction maps to automatically annotate protein secondary structure elements as well as RNA/DNA. It can be straightforwardly applied to annotate newly reconstructed maps to support domain placement or to supply a starting point for main-chain placement. Due to its high recall and precision rates of 95.1% and 80.3%, respectively, on an independent test set of 122 maps, it can also be used for validation during model building. The trained network will be available as part of the CCP-EM suite.

## Introduction

In recent years, three-dimensional density maps reconstructed from single particle images obtained by electron cryo-microscopy (Cryo-EM) have reached unprecedented resolution. These structures allow us to identify new drug targets, for example in the Zika virus^1^, to fight tuberculosis^2^ or to understand the fundamental processes of life, such as the process of translation through ribosomes^3^. However, modelling an atomic structure to these maps remains difficult as researchers mostly rely on algorithms developed for the interpretation of crystallographic electron density maps. In X-ray crystallography, the measured diffraction corresponds to the amplitudes of the Fourier transform of electron density, as the X-rays interact with the electrons in the molecular assemblies in a crystal and the phases are reconstructed only during refinement. In electron cryo-microscopy, on the other hand, the measured micrographs already contain phase information, but are very noisy, which is overcome by 3D-reconstruction and averaging. The individual micrographs show the interaction of the electron beam with the entire electrostatic potential of a single molecular assembly. Hence, Cryo-EM reconstruction maps differ in both their nature and error distribution^4–6^ from crystallographic electron density maps. Consequently, their modelling might be improved greatly by tools that take into account these specific properties of the data at hand. Such modelling tools should not only provide good functionality, but also be easy to use and freely available to academic users worldwide.

Parallel to the advances in Cryo-EM during the last decade, deep neural networks achieved remarkable image segmentation capabilities^7^, making them the most powerful machine learning approach currently available. Convolutional neural networks (CNN) combine traditional image analysis with machine learning by cascading layers of trainable convolution filters and are exceptionally well suited for volume annotation. They have been successfully applied to biological problems such as breast cancer mitosis recognition^8^ and, in conjunction with encoder-decoder architectures, to volumetric data segmentation^9,10^. Given that a Cryo-EM reconstruction map is essentially a three-dimensional image volume, CNNs seem a good choice for their annotation if good ‘ground truth’ data to train the network could be provided.

In this work, we demonstrate that deep neural networks are not only capable of annotating protein secondary structure, but also oligonucleotides (RNA/DNA) in Cryo-EM maps, and provide a pre-trained network, named *Haruspex*. Assigning a fold to regions in a Cryo-EM map is the first step in modelling a structural region. This can be a major challenge, in particular for novice users, in low resolution regions, or when little is known about the composition of the macromolecular complex in question. Haruspex can be readily used to annotate Cryo-EM maps, which will prove useful to facilitate model building and to support the placement of known domain folds, thus accelerating the modeling process and improving the accuracy of Cryo-EM derived molecular structures.

## Results

### Network architecture and implementation

In low-resolution Cryo-EM maps, α-helices can often be discerned as long cylindrical elements. This has been exploited by the program *helixhunter*^11^, which searches for prototype helices in reconstruction maps using a cross-correlation strategy. β-Strands are more difficult to identify as they are more variable in shape and therefore require morphological analysis^12^. A combination of these approaches led to the development of *SSEHunter*^13^, which uses a density skeleton to detect secondary structures. Deep learning offers an alternative approach: Fully convolutional networks^9,14^ allow a swift generation of segmentation maps for objects of variable size. Here, we employ a state of the art U-Net-style architecture^9^ to demonstrate that at an average map resolution of 4 Å or better, experimentally derived reconstruction maps allow training of a well-performing network that can be used for a wide range of specimens - with no re-training being necessary. The network was implemented with Tensor Flow^15^ and processes 40^3^ voxel segments with a voxel size of 1.0-1.2 Å^3^ (covering a secondary structure element and its immediate surroundings) to annotate 20^3^ voxel cubes (corresponding to the center of the input volume). The output volume has four channels containing the probabilities that the voxel is part of an α−helical or β-strand protein secondary structure element, nucleotide, or unassigned. 40^3^ voxel segments were chosen as a compromise between computational power and network complexity on one hand and covering the secondary structure including surrounding interaction partners on the other hand; A 40^3^ voxel segment covers 40-48 Å^3^; 
an average α-helix with 10 residues, for example, is 15 Å in length^16^.

The input is a single channel containing the reconstruction density. During prediction, this three-dimensional volume is passed through multiple convolutional layers (image filters) that extract learned image features relevant for structure detection, and through pooling layers, which select the most significant of the detected features. In the second (‘upconvolutional’) part of the network, these activations are combined with higher-level activations of the network to recover spatial detail. The output has four channels representing the probabilities for the four classes (helix, sheet, nucleotide, unassigned) and represents the annotation of the central 20^3^ voxel cube of the input volume.

### Training data selection

For network training, we pre-selected EMDB (Electron Microscopy Data Bank^17^) reconstruction maps with an average resolution of 4 Å or better as stated in the EMDB entry. From 576 entries (as of 15/2/2018), we picked 293 EMDB/PDB (Protein Data Bank^18^) pairs by three criteria: (1) map and model represent the same structure and fit visually well to each other; (2) presence of at least one annotated α−helix or β-sheet in the PDB model; (3) preference of higher resolution maps in case the same authors deposited several instances of the same macromolecular complex (as the model was most likely fitted to the highest resolution map). Maps with severe misfits, misalignments, or models without corresponding reconstruction density (and vice versa) were omitted. Visual evaluation was supplemented with a comparison between the entire map and the part which is occupied by the model using histograms, mean and median values; this provided an additional test of how well map and model fit each other. Furthermore, the training data were filtered by map root mean square deviations (r.m.s.d.) values (see below).

### Training data annotation

To generate ground truth data for network training, a python script was implemented to automatically annotate the reconstruction map according to the deposited structural model as α−helical, β-strand, nucleotide or unassigned. The script extracts the original annotations from PDBML format^19^ files using a custom parser. To obtain suitable training data, additional secondary structure information was necessary. We implemented a variant of the DSSP algorithm^20^ omitting strand direction, and a torsion angle based secondary structure detection inspired by STRIDE^21^: annotated or DSSP-detected secondary structures were extended by neighboring amino acids, if they matched the same Ramachandran profile. Before usage, the voxel size of the reconstruction map was re-scaled to 1.1 Å if outside [1.0; 1.2] Å.

Following that, if a secondary structure was identified, and if the average main chain atom map r.m.s.d. (root mean square of the map density distribution) was above 2, all voxels within 3 Å of backbone atoms were annotated accordingly. Secondary structure residues below 2 but ≥ 1.0 r.m.s.d. were masked and excluded from error calculation during training. All voxels not within 5 Å of model atoms, but with density ≥ 1 r.m.s.d. were masked and excluded from training, as they had high density, but were not modelled. The remaining voxels were marked as ‘unassigned’.

### Network training

The maps were split into a total of 2183 segments of 70^3^ voxels, of which 110 segments (5%) were set aside for evaluation during network training. Each segment had to contain at least 100 atoms ≥ 1.0 r.m.s.d., a backbone mean density of ≥ 3 r.m.s.d., and at least 5% of the total segment volume annotated. The training data were augmented through on-GPU 90° rotations (24 possibilities), and by randomly selecting a 40^3^ voxel sub-segment (translational augmentation). Furthermore, false negatives were weighted 16-fold stronger than true negatives. This was necessary as the majority of the space within a reconstruction map is not made up of secondary structure associated voxels and thus the network can reach ∼70-90% accuracy by sticking to ‘unassigned’ (not α-helical, β-sheet or oligonucleotide) secondary structure) only. The 16-fold weighting factor for false negatives was established empirically. Further optimization of weighting might improve the false positive rate in the future. The network was trained for 40,000 steps with 100 segments employed per step.

### Network performance test

After training, the network was tested on an independent set of 122 EMDB maps (selected by the same criteria as training data and deposited after February 2018). For evaluation, we investigated residues with mean backbone densities ≥ 1.0 r.m.s.d. and compared the predicted secondary structure on a per-residue basis with the one derived from the deposited PDB model. For this analysis, the r.m.s.d. value given in the header of each map file was used. Using this criterion, the network achieved similar performance on training, evaluation and test data. Over all test maps, there were 75.4% true positives *r*_*p*_ (correctly predicted residues), 18.8% false positives f_p_ (wrongly predicted residues) and 4.0% false negatives *f*_*n*_ (non-predicted residues), resulting in a median recall rate 100\**r*_*p*_(*r*_*p*_ + *f*_*n*_)^-1^ of 95.1% and a precision 100\**r*_*p*_(*r*_*p*_ + *f*_*p*_)^-1^ of 80.3%. Precision and recall did not correlate significantly with average resolution (as given in the EMDB entry), Molprobity^22^ score or deposition date.

The corresponding residue-level F_1_ score (harmonic mean of precision and recall) on the test set for Haruspex (87.05%) is the highest reported so far on the per-residue-level when compared to other recent work (Li 2016, Haslam 2018 and Subramaniya 2019). Direct comparison of these values is, however, difficult since these other networks were tested on small test sets of lower-resolution simulated and experimental maps, whereas we used a large set of exclusively experimentally derived higher-resolution maps. Moreover, these networks did not annotate oligonucleotides, which affects the composition of the F_1_ score. In a recent preprint ^23^, the authors use deep learning for atom-level prediction and report 88.5% correctly predicted C_α_ atoms on 50 pre-cleaned experimental maps at 4.4 Å or better, which suggests similar performance for their intermediate secondary structure prediction.

As a typical example, human ribonuclease P holoenzyme (EMDB entry 9627) illustrates the power of our approach (see Fig. 1). Haruspex is not only able to accurately predict the RNA vs. protein distribution in this complex, but also correctly assigns secondary structure elements in the protein areas with only a few exceptions. These notably include the turn of the RNA (upper left of the structure), regions that resemble β-sheets but do not follow the characteristic hydrogen bonding pattern, as well as secondary structure elements currently not covered by Haruspex, such as polyproline type II (P_II_) helices (Fig.2C and D). Additional examples are shown in Fig 3.

**Figure 1.**
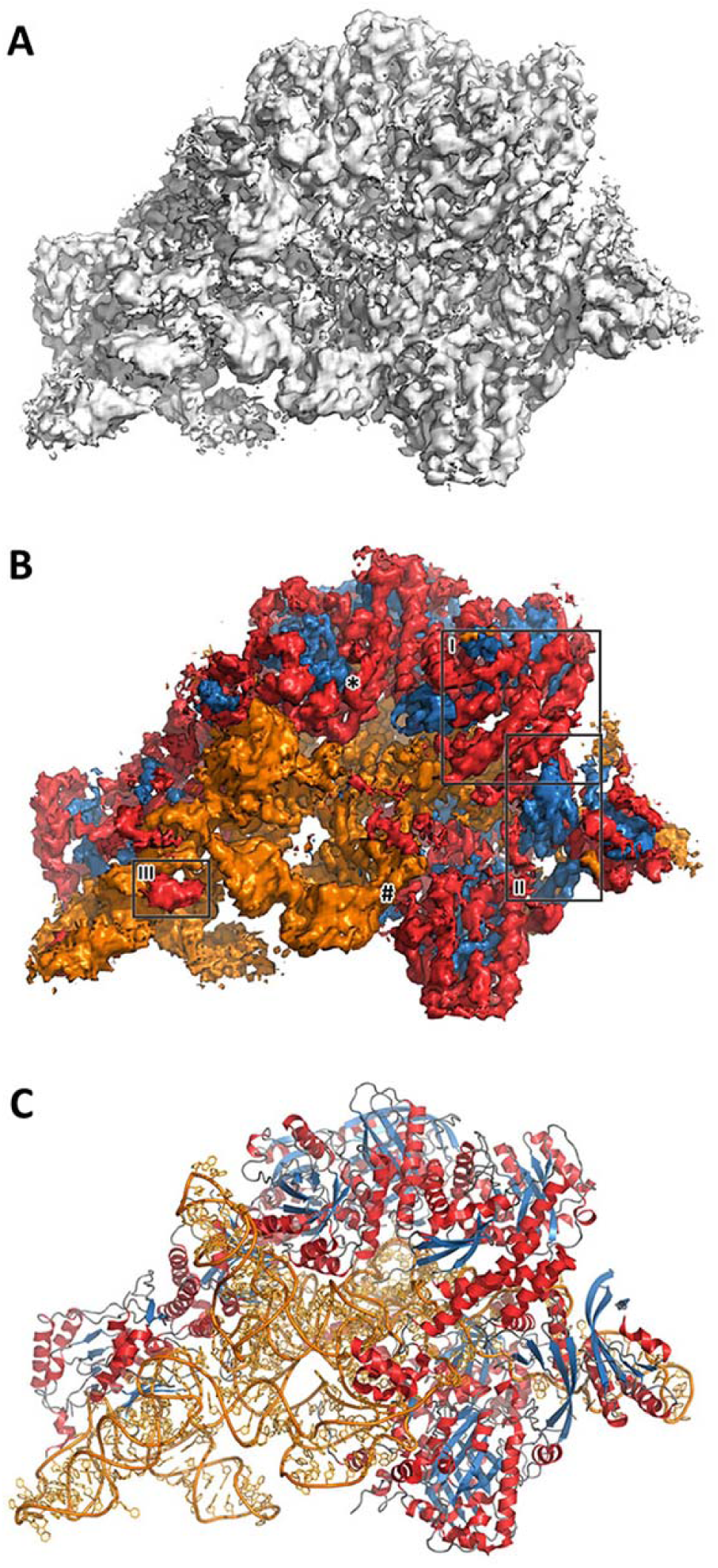
Typical example of Haruspex annotation. **A.** Reconstruction map for the human Ribonuclease P holoenzyme (EMDB entry 9627). Manual assignment of secondary structure features can be difficult, in particular if the composition of a macromolecular complex is unknown. The shown surface corresponds to an r.m.s.d. of 0.04 with no carving. **B.** Secondary structure as identified by our network in the map, is projected onto the surface. Orange corresponds to RNA/DNA; red to helices and blue to sheets. This was a fairly typical test case with 70.5% true positives, 18.8% false positives and 10.7% false negatives. Recall was 86.8% and precision 79.0%. Region (I) depicts a well-predicted α-helical structure, (II) a β-sheet and (III) RNA misinterpreted as an α-helix. **C.** The deposited model PDB 6AHU for this map is shown in comparison. The regions depicted in Fig. 2C and 2D are marked **#** and **\***, respectively.

**Figure 2.**
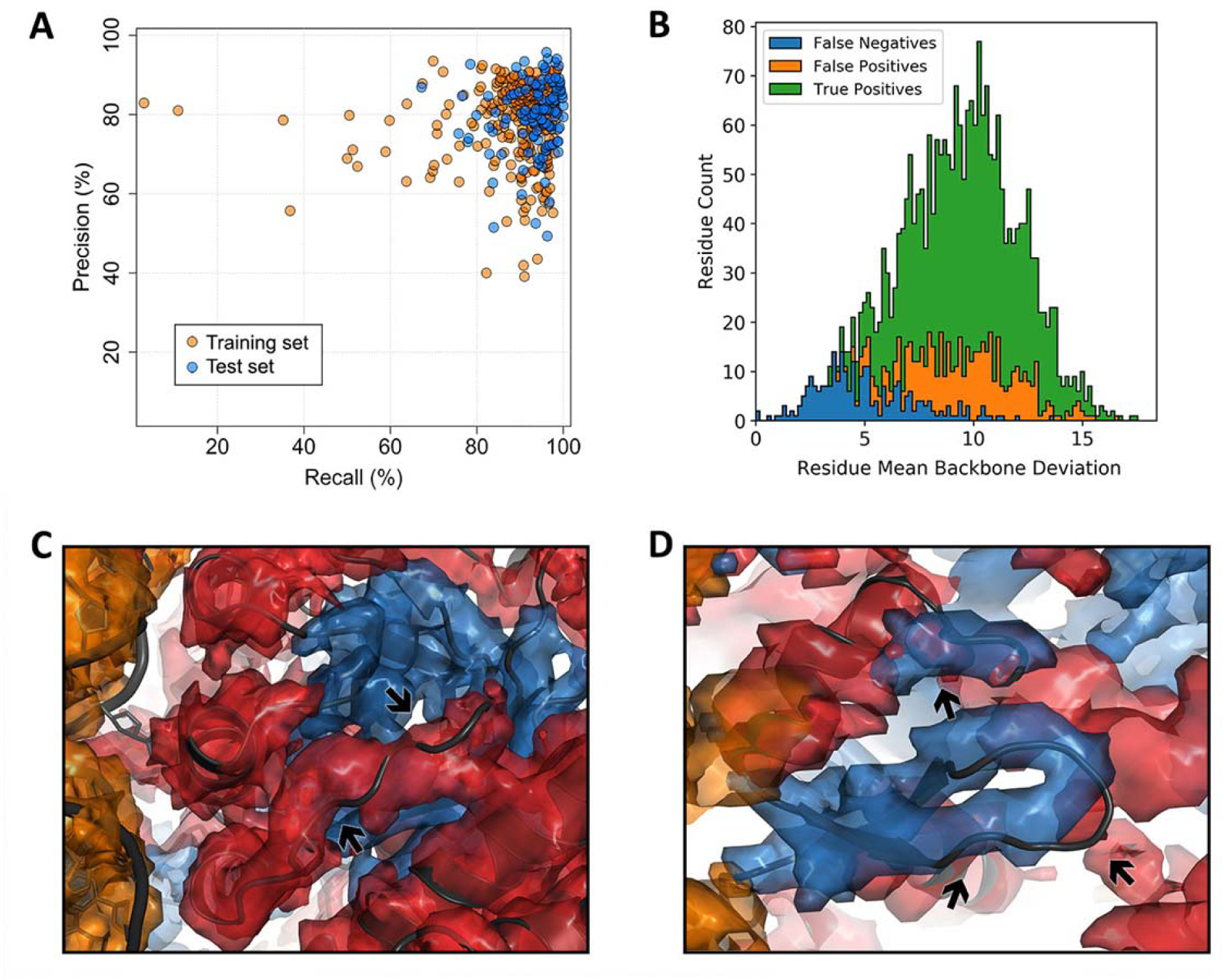
Network performance. **A.** Network precision vs. recall rates, with one marker per EMDB entry (training set entries are shown as orange, test set entries as blue markers). Both perform similarly well, with the training set producing a few more outliers. **B.** Frequency vs. map r.m.s.d. level for EMDB 9627 on a per-residue basis: True positives (green), false positives (orange) and false negatives (blue). This plot is typical: false negatives often occur in low density map regions. **C.** α-Helical false positives (PDB 6AHU, J131 – 139): The model partly occupies the conformational space of a polyproline type II helix (P_II_), which is often misinterpreted as α-helical and may have been modelled incorrectly (given that the model does not completely fit the density). **D.** False positives in a β-sheet (6AHU, B215-B221). The deposited model does not maintain the hydrogen bonding that defines a regular β-sheet; to the network, however, the fold still ‘looks’ like a β-sheet to the network and a third segment (top) is also assumed to be part of it.

**Figure 3.**
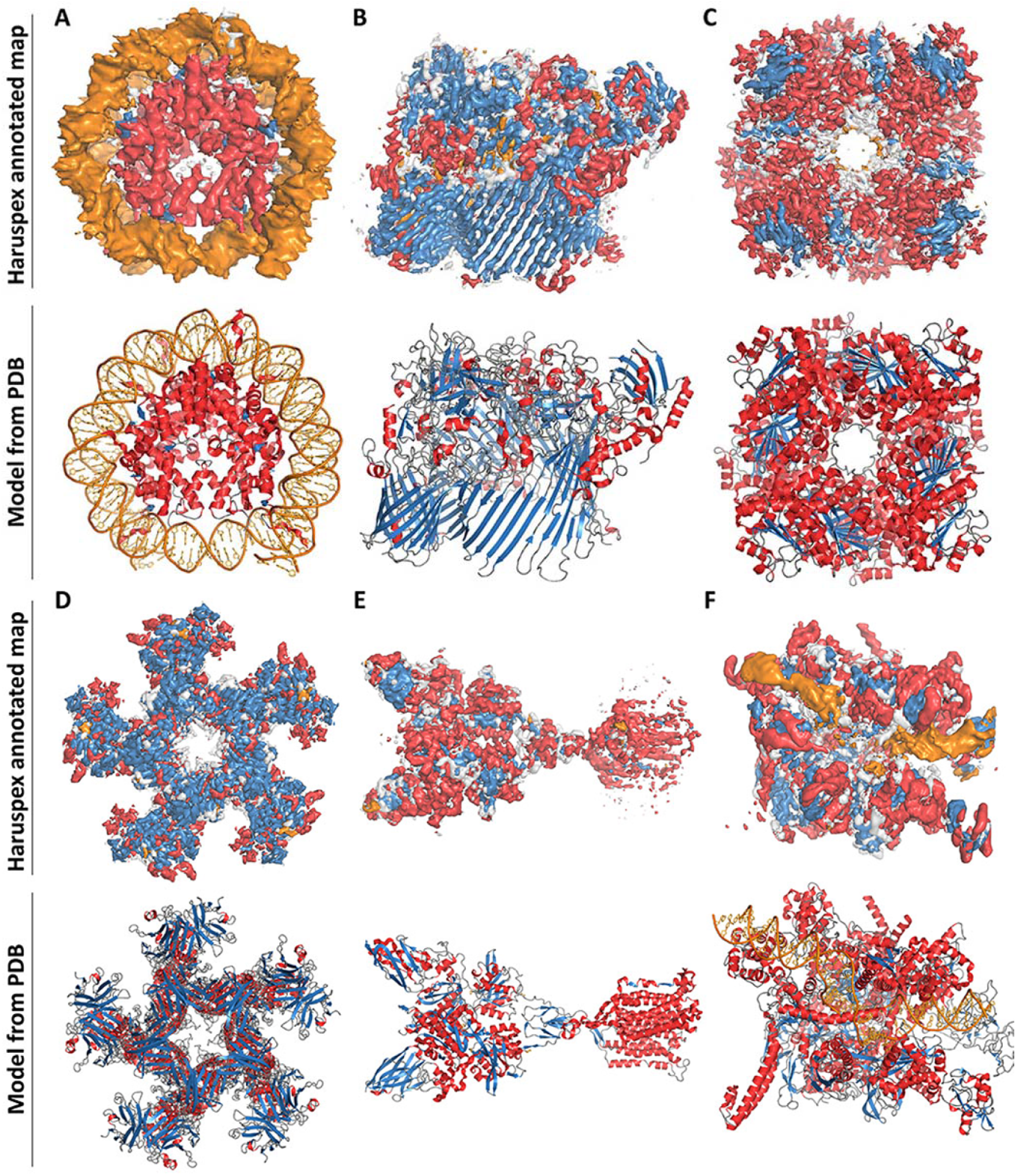
Additional examples from the test set. Top: Annotated map. Bottom: Deposited structure for comparison. Orange corresponds to RNA/DNA; red to helices; blue to sheets and grey regions were not assigned any secondary structure. **A.** Nucleosome from *Xenopus laevis*, average map resolution 3.8 Å (map: EMDB 4297/model: PDB 6FQ5): recall 98.5%, precision 94.0% **B.** *Flavobacterium johnsoniae* Type 9 protein translocon, average map resolution 3.5 Å *(map: EMDB 0133 /model: PDB 6H3I): recall 96.3%, precision 49.3%* **C.** Leucine dehydrogenase from *Geobacillus stearothermophilus*, average map resolution 3.0 Å *(map: EMDB 9590 /model: PDB 6ACF): recall 89.8%, precision 85.7%* **D.** *Escherichia coli* Type VI secretion system, average map resolution 4.0 Å *(map: EMDB 9747/model:PDB 6IXH): recall 95.9%, precision 70.9%* **E.** Homo sapiens metabotropic glutamate receptor 5, average map resolution 4.0 Å *(map: EMDB 0345/model: PDB 6N51): recall 95.9%, precision 71.7%* **F.** Bacterial RNA polymerase-sigma54 holoenzyme transcription open complex, average map resolution 3.4 Å *(map: EMDB 0001 /model: PDB 6GH5): recall 94.2%, precision 67.5%.*

### Haruspex Usage

Haruspex can be used as a command line tool, which reads in an MRC format reconstruction map. No further parameters are needed and a prediction for a single map takes approximately 30 seconds to a few minutes on a normal workstation, depending on the available hardware (it can be used with or without GPU); on an older laptop, the annotation may take as long as 45 minutes for a large structure. The output consists of four MRC format maps corresponding to the α−helical and β-strand protein, nucleotide, and ‘unassigned’ portion of the input map. These maps can be displayed in *Coot*^24^, *Pymol*^25^ or *Chimera*^26^ and together represent the entire input map. The source code and trained network, including Docker containers, can be downloaded from https://github.com/thorn-lab/haruspex, and will soon be available as part of *CCP-EM*^27^.

## Discussion

### Network Performance

Here we have described the development of the neural network Haruspex for the annotation of protein secondary structure and RNA/DNA in Cryo-EM reconstruction maps in order to facilitate the modelling of such maps. We trained Haruspex on 293 experimentally derived reconstruction maps at 4 Å or better and obtained recall and precision rates of 95.1% and 80.3%, respectively, on an independent test set of 122 maps. The pre-trained network can be readily applied to annotate newly reconstructed maps to support domain placement or to supply a starting point for main-chain placement.

When considering the 18.8% false positives and 4.0% false negatives, two fundamental limitations in the annotation of EMDB maps should be considered: firstly, the map can be wrongly modelled (see Fig. 2C), which biases our annotation towards human modelling errors. Secondly, the deposited model may have been built employing additional information, such as structure-specific information from an external source, for example backbone folds established prior by crystallographic means^28^, NMR or structure prediction - or more than one map generated from different particle alignments^29^. This would in particular introduce higher rates of false negatives at the outer edges of the map, where the model covers secondary structure that was established by other means, but the map does not provide enough information to make this assignment.

Closer inspection reveals that false positives are often elements closely resembling helices, sheets or RNA/DNA (see Figs. 2 and 3). In particular, semi-helical structures, β-hairpin turns and residues belonging to polyproline type II (P_II_) helices^30^ are misclassified as α-helical - and loosely parallel structures without the typical hydrogen-bond pattern are frequently misclassified as β-strands. In the case of P_II_ helices, this is partly due to the STRIDE-like annotation. It would be very desirable to quantify the false positives in this respect, but this was not possible within the scope of this work, as no automatic annotation algorithms seems to exist for such cases. For the future development of Haruspex, predicting additional classes, such as β-turns, poly-proline helices and perhaps even membrane detergent regions would be very desirable, as this would potentially lower the number of incorrectly identified secondary structure elements, while at the same time supplying additional information to users.

### Resolution range and comparison to similar algorithms

Haruspex was trained for average resolutions as low as 4 Å, and at the current rate by which the resolution of published Cryo-EM maps increases, the average resolution in published maps may well be 3.5 Å by 2021 (P. Emsley, personal communication, April 2019). Irrespective of this, we will extend our approach to lower resolution data in the future, where our automated annotations should be even more advantageous for users. Still, low resolution experimental maps with a well-matching model for training and testing such a network are difficult to obtain. This obstacle has previously been faced by Si et al.^31^ (*SSELearner*), Li et al.^32^ and Subramaniya et al.^33^ (*Emap2Sec*) who developed machine learning approaches for protein secondary structure prediction in Cryo-EM maps, but not oligonucleotides^32^, and consequently resorted partly to simulated maps generated with *pdb2mrc*^34^. These simulated maps lack the error structure and processing artefacts found in experimentally derived reconstruction densities^4–6^, as they assume a perfectly processed data set of a homogenous sample where all atoms interact with the electron beam as if they were uncharged and unbound. Si et al. tested their support vector machine on 10 simulated maps of relatively small structures (< 40 kDa) and, as available data were still very limited in 2012, only 13 experimental maps paired with individually selected training maps. Haslam et al. ^35^ used a 3D U-Net, which was trained on 25 simulated and 42 experimental maps between 3-9 Å resolution to predict helices and sheets obtaining an *F*_*1*_ score 2(recall^-1^ + precision^-1^)^-1^ between 0.79 and 0.88. However, the network was only tested on six simulated maps and one experimentally derived map. We, on the other hand, used a total of 293 experimentally derived maps in a semi-automated workflow to provide a more realistic training environment. Furthermore, the amount of newly released high resolution structures in conjunction with our processing infrastructure permitted us to test our network performance on a representative set of 122 unique depositions. The semi-automated workflow for the selection and annotation of training data (see Methods) allows for an easy expansion of ground truth data and re-training. However, given that Haruspex has already been trained on a diverse range of macromolecular structures, the network can be used to interpret any map at 4 Å or better without any additional (re-)training necessary. The newest version of Haruspex will regularly be distributed as part of *CCP-EM*^27^.

### Augmentation of automatic model building

Haruspex ideally complements tools for automatic map-based structure building such as MAINMAST^36^, RosettaES^37^, ARP/wARP^38^, phenix.map_to_model^39^ or Buccaneer^40^ by providing an independent method to locate secondary structure elements of proteins to assist the validation of an automatically built protein main-chain. Haruspex may even be employed in the future to serve as starting point for such methods. The ability of Haruspex to automatically recognize RNA/DNA is of particular interest for the analysis of ribosomes, spliceosomes and polymerases which all contain substantial amounts of oligonucleotides. As these and similar structures are among the most common specimens studied by single-particle Cryo-EM, Haruspex, which, to our knowledge, is the first to use machine learning for the identification of nucleotides in Cryo-EM reconstruction maps, offers a unique advantage for the analyses of these structures.

### Summary

We show that a neural network can be used to automatically distinguish between nucleic acids and protein and to assign the two main protein secondary structure elements in experimentally derived Cryo-EM maps. This technique will render the process of protein structure determination faster and easier. Haruspex was trained on a carefully curated ground truth dataset based entirely on experimental data from EMDB. The pre-trained network can be straightforwardly applied to annotate newly reconstructed Cryo-EM density maps. Besides guidance for domain placements, the network also proves useful for model validation during building due to its high median recall and precision rates of 95.1% and 80.3%, respectively, as has been demonstrated by early users at our institute, for example in the modelling of the mycobacterial type VII secretion system^2^. We plan to refine and adapt the network as new data become available, extend the approach to lower resolution and more structural classes in the future.

## Methods

### Training Data

We queried the Electron Microscopy Data Bank (EMDB) for all single particle Cryo-EM maps with a resolution ≤4 Å, for which corresponding protein models were available in the Protein Data Bank in Europe (PDBe), yielding 576 map and model pairs as of February 2018. We filtered these EMDB/PDB pairs by the following three criteria: (1) Visual good fit between map and model; (2) presence of at least one annotated α-helix or β-sheet; and (3) preference of the highest resolution map in case the same authors deposited several instances of the same macromolecular complex. Maps with severe misfits, misalignments, or models without corresponding reconstruction densities, and vice versa, were discarded. After applying these criteria, we retained 293 map/model pairs for generating the training data.

To extract secondary structure information from the PDB data, we developed a custom parser for the PDBML^19^ format based on xmltodict^41^. To obtain additional secondary structure information, we implemented a variant of the DSSP algorithm^20^ without strand direction, and a torsion angle based secondary structure detection inspired by STRIDE^21^: annotated or DSSP-detected secondary structures were extended by neighboring amino acids if they matched the same Ramachandran profile.

### Annotation of reconstruction maps

For every entry pair, the augmented model was then superimposed on the map and all voxels within 3 Å of a protein backbone atom, or, in the case of nucleotides, within 3 Å of any atom, were assigned to the respective class (helix, sheet or nucleotide) if their value was higher than ½ of the average backbone density of the helix, sheet or nucleotide in question. Secondary structures with a backbone standard deviation of <2 σ and atoms without secondary structure assignment were either excluded, as they were likely incorrectly modelled, misfitted, or flexible structures, or labelled as ‘unassigned’. For some training data pairs, such as virus capsids, only small or partial protein models were deposited for large Cryo-EM maps, resulting in well-defined high-density regions without model coverage. These regions would not get annotated and hence result in false positives if the network tried to predict the actual structure. To mitigate this, all voxels with density ≥ 1.0 r.m.s.d. but not within 5 Å of a model atom with density ≥ 1.0 r.m.s.d. were masked as unmodeled density and hence did not contribute to training.

Since our network generated a single class label as output, the reconstruction density of the secondary structures must be converted to a strict assignment to one of the three classes in order to be used as training examples. For each secondary structure, the reconstruction map density was multiplied by the backbone standard deviation and rescaled to an output density between zero and one (corresponding to 0.5 and 1.0 times the average backbone density of the local secondary structure element) for each label type. The highest channel value determined the voxel class. If multiple channels shared the same value, sheets took precedence over oligonucleotides, which took precedence over helices. Voxels where all channel values were below 0.01 were assigned the ‘unassigned’ class. Finally, reconstruction maps were rescaled to a voxel size of 1.1 Å if they were outside of [1.0; 1.2] Å.

### Generation of training segments

To generate the 70^3^ voxel sized segments needed for training, candidate volumes were sampled from the entire map, and segments with a mean backbone density < 3.0 r.m.s.d., less than 5% annotated volume, or less than 100 atoms with standard deviation ≥ 1.0 r.m.s.d. were discarded. This resulted in altogether 2183 training segments, of which 110 segments (5%) were held back for evaluation during training. To generate additional segments for training, we applied rotations in steps of 90° around all three axes, resulting in 24 rotated versions of each segment that could all be used as separate training volumes since the convolutional network is not rotation-invariant. Segments were further augmented during training by using a randomly translated 40^3^ sub-cube for each step.

### Network Architecture

We used a state-of-the-art U-Net-like encoder-decoder architecture^9,10^ (see Fig. 4) with a single input channel (the reconstruction density). This architecture is a variant of so-called fully convolutional networks where spatial information and object details are encoded, reduced by pooling layers and then recovered again with up-sampling or transpose convolutions; the term U-Net arises from the U-like shape of the data flow. The encoding branch consisted of two 3×3×3 convolutional layers with 32 and 64 feature channels, respectively, followed by max-pooling layers. Another convolutional layer with 128 feature channels followed by a max-pooling layer finally resulted in an 8^3^ cube with 128 feature channels at the deepest layer of the network. This cube was passed through another convolutional layer with the same data padding in order to preserve its dimensions. A fully connected layer was considered, but not chosen due to its high memory and performance cost. The decoding branch of the U-Net was made of two blocks, each consisting of a deconvolution followed by two 3×3×3 convolutions (128 feature channels in the first, 64 and 32 channels in the second block to restore symmetry) with concatenated sections of the corresponding layer in the encoding part. The output part consists of a final 1×1×1 convolution followed by a soft-max output layer. The output layer reproduced the central 20^3^ voxel cube of the input layer in four annotation channels representing co-dependent probabilities for the four classes (helix, sheet, nucleotide, unassigned) summing up to one. The highest channel value determined the predicted class. Implementation was realized using TensorFlow ^15^. The network was trained end-to-end by comparing the predicted class of each voxel to the annotated EMDB model using cross-entropy loss, back propagating the error through the network, and adapting the network weights to iteratively minimize the error.

**Figure 4.**
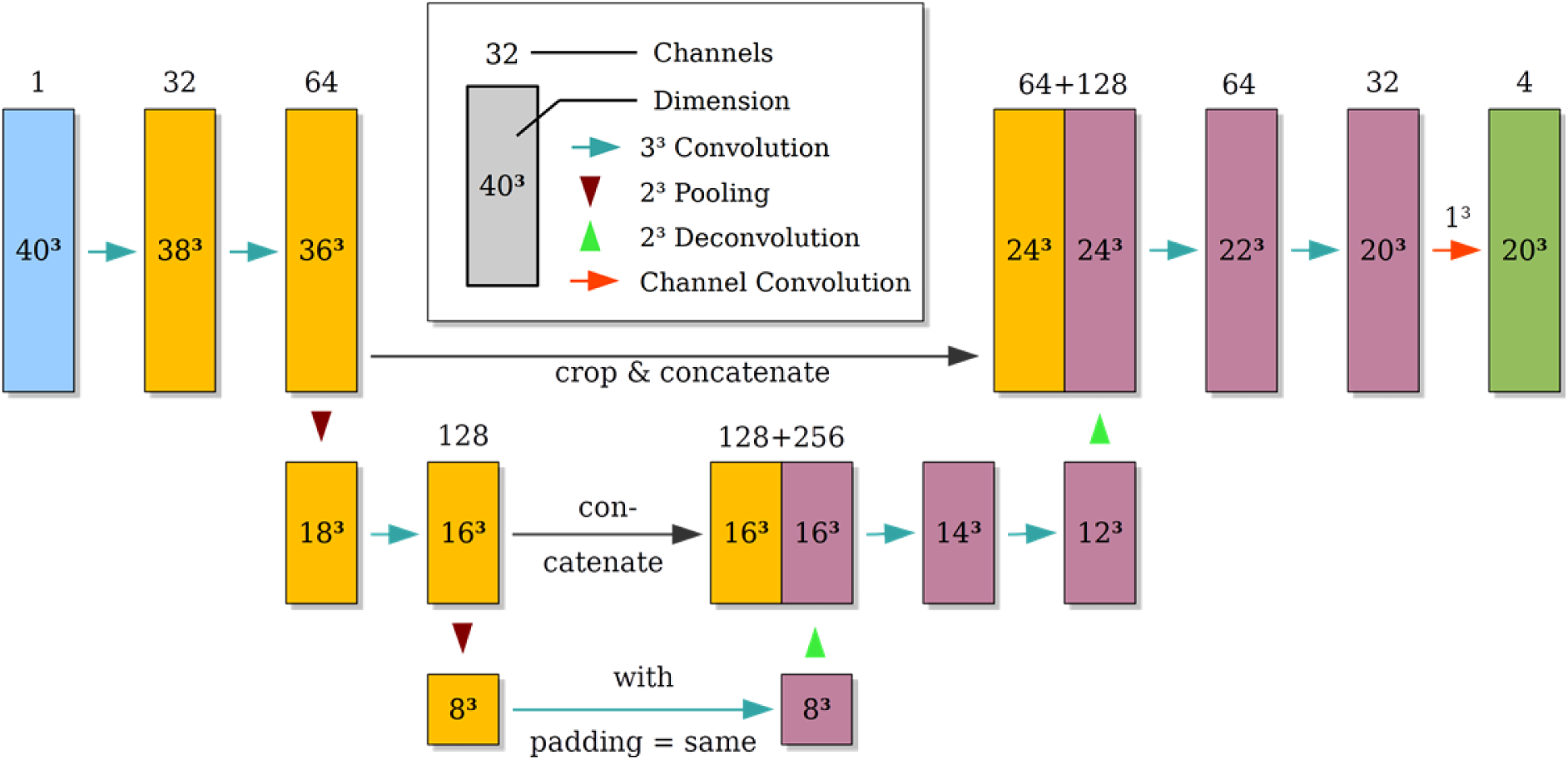
Haruspex neural network architecture. The network consists of multiple interconnected layers, shown as rectangular boxes. The layers are connected by convolution and pooling operations (arrows). Layer height represents the level of abstraction: lower layer data, generated by pooling operations, contain more abstract representations of the map. Input data (blue) is fed into the downconvolutional arm (yellow) in order to extract valuable information, which is then combined with previously discarded information through concatenations in the upconvolutional arm (purple) to compute annotated output data (green) for a subsection (20^3^) of the input volume (40^3^). Our network consists of two encoder blocks, containing altogether three convolutional layers (3×3×3) and two pooling layers. This is followed by two decoder blocks, one with upconvolution followed by two 3×3×3 convolutions and 128 feature channels, and one with upconvolution followed by two 3×3×3 convolutions with 64 and 32 feature channels, with concatenated sections of the corresponding layer in the encoding part. The output part consists of a final 1×1×1 convolution followed by a soft-max output layer. This results in 13 layers in total (12 + 1 convolution at bottom).

### Network Training

The network was trained for 40,000 steps on training batches of 100 random segment pairs per step corresponding to 80 epochs, using ADAM stochastic optimization^42^ with a learning rate of 0.001, β_1_ = 0.9, β_2_ = 0.999 and ε = 0.1. Error assignment for backpropagation was performed using cross-entropy loss, where the target class was represented in one-hot encoded binary format (1 for the target class, 0 for the other three classes). To account for class imbalance, voxels were weighted according to overall class occurrence in the training data. Furthermore, non-true negatives were weighted 16-fold stronger than true negatives

## Acknowledgements

This work was supported by the DFG (project TH2135/2-1), the High Performance Computing Cloud of Würzburg University (DFG project 327497565) and the Rudolf Virchow Center for Experimental Biomedicine. We would like to thank Bettina Böttcher, Niko Grigorieff, Jane Richardson, Christopher Williams, Tom Burnley and Jola Mirecka for fruitful discussions; Virginie Uhlmann and the image analysis journal club for helpful comments on the preprint; and Bernhard Fröhlich for great computational support.

